# A new species of the genus *Carlogonus* (Spirostreptida: Harpagophoridae) from West Bengal, India

**DOI:** 10.1101/2021.04.25.441382

**Authors:** Somnath Bhakat

## Abstract

A new species of *Carlogonus, Carlogonus bengalensis* is described from West Bengal, India. The adult is blackish brown in colour with a yellowish curved tail, round body with 60 segments, 55 mm in length, 5^th^ segment of male bears a hump, telopodite of the gonopod long, flat and band like with a single curved antetorsal process, mesal process with a red spine, proplical lobe with a curved orange spine, inner surface of metaplical fold with sigilla, palette spatula like and a few blepharochatae at the apical margin. Male bears white pad on femur and tibia. Comparison was made with the “*exaratus* group” of the genus *Carlogonus*.

## Introduction

A few reports mostly on different aspects of millipedes (except taxonomy) especially on Polydesmid millipedes are available from West Bengal (India). Except the study of Bhakat (2014), there is no report on any aspect of Spirostreptid millipede from this region. Of the Spirostreptid millipede, genus *Carlogonus* Demange, 1961 includes three Harpagophorid millipede species described in the Southeast Asian genus *Thyropygus* Pocock, 1894 and the South Asian genus *Harpurostreptus* Attems, 1936. Distribution of this genus is restricted to India and Srilanka only. Till date 10 species are reported of which nine from India and one from Sri Lanka (De Zoysa et al., 2016; Golovatch and Wesner, 2016; Sankaran and Sebastian, 2020). Except *C. exaratus*, all the Indian species of *Carlogonus* are geographically restricted to southern part of India (Demange, 1961, 1977a, b, 1981, 1983; Mauries et al. 2001; Golovatch and Wesner, 2016; Sankaran and Sebastian, 2020).

I have collected a huge number of different species of millipedes from Birbhum (a district of West Bengal) in the period 1983-1993 and preserved the specimens in 4% formaldehyde solution in my own custody. In that period, I send a few specimens to Dr. R. L. Hoffman, Virginia Museum of Natural History, USA, for identification. Among those, he identified one of the specimens as genus *Carlogonus*. In this paper, I described that species as *Carlogonus bengalensis* sp. nov.

## Materials and methods

Millipedes were hand collected from the road side and from the litter in the month of June, 1992. Collections were made at dawn (4.30 a. m. to 5.30 a. m.) as they are nocturnal in habit. Hand collected specimens were euthenised by cooling and then preserved in 4% formaldehyde. Specimens were examined using simple and stereoscopic microscopes. In some cases sketches were drawn with the help of camera lucida. Photographs were taken from the microscope by camera. Morphological measurements were made with the help of digital slide calipers and an ocular micrometer.

Here I followed the modern terminology of gonopod for description of the species (Hoffman, 2008). The following old terms are replaced by the new terminology: Anterior coxal fold= proplica, posterior coxal fold= metaplica, tibial spine= antetorsal process, gonocoel= gonochiasma.

## Result

### Systematics

Order: Spirostreptida Brandt, 1833
Suborder: Spirostreptidea Brandt, 1833
Family: Harpagophoridae Attems, 1909
Subfamily: Harpagophorinae Attems, 1909
Tribe: Harpurostreptini Hoffman, 1980
Genus: *Carlogonus* Demange, 1961

#### Diagnosis

Comparatively large sized millipede with a long, pointed, curved anal spine turned downwards. Gonopodal sternite is short with round posterior parts. Anterior coxal fold is lamelliform while posterior with internal lateral lobes. Anterior coxal leaves bear hooks and appendages. A hook or a long pointed appendix is present on the external lateral side of each leaf. Telopodite long, flattened, circular or spiral in shape and with femoral and tibial spine. Palette unbranched with blepharochaetae (Attems, 1936; Demange, 1977a, b; Bano, 1998).

Genus *Carlogonus* differs from its congeners by one or more distinguishing characters. The genus can be distinguished from *Harpurostreptus* by the presence of a voluminous tibial spine (vs. no tibial spine), and without spine along the entire length of the inner side of spiral telopodite (vs. a number of spines all along its inner side). In *Gamognathus*, telopodite long or short and divided into 2-3 branches at its tip while in *Calogonus* it is long and with a single branch. In *Carlogonus*, palette of telopodite unbranched but in *Gamognathus* it may be branched or unbranched. In *Janardananeptus*, gonopod sternite is long and telopodite with branched palette (vs. short sternite and unbranched palette in *Carlogonus*). *Carlogonus* is distinguished from *Organognathus* by the presence of long, flattened, somewhat spiraled telopodite with unbranched palette and by the absence of spine like apophyse on the posterior coxal fold vs. short, not rolled into a spiral telopodite with branched palette and have strong spine like apophyse on the posterior coxal fold (Verhoeff, 1936; Attems, 1936; Carl, 1941; Demange, 1961; Bano, 1998)

***Carlogonus bengalensis* sp. nov.** (Fig. 1)

**Fig. 1.**
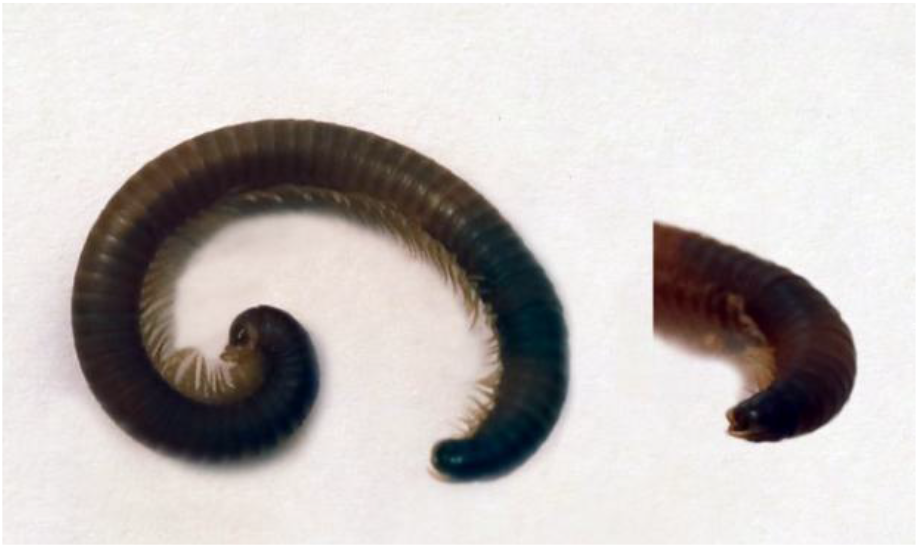
Male *Carlogonus bengalensis* sp. nov (the posterior portion is shown seperately).

### Type material

Holotype: Male (length 55 mm, width 4.0 mm, 60 segments, 218 legs, ommatidia 11 rows); Suri (87°32’00”E, 23°55’00”N), Birbhum district, West Bengal, India; 27. VI. 1992; S. Bhakat; Zoological Museum, Dept. of Zoology, Rampurhat College, Rampurhat-731224, Dist. Birbhum, West Bengal, India and in the personal collection of the author.

Paratype: One male and one female; same spot on 29. VI. 1993. Other information are same as holotype.

#### Etymology

The specific epithet refers to the name of the state “Bengal” (West Bengal) from where the new species belongs.

#### Description

Male: length 55.4mm, width 4.0mm. (n= 2). Female: length 57.2mm, width 4.9mm.

Head: Smooth with sparsely distributed small setae on the margin, no interocular line. Antenna of male is comparatively longer (3.52mm vs. 2.92mm) and 0.88 of body diameter. The terminal article of antennae is smallest and bears four apical cones. Length ratio of antennomeres in male: 1.43: 1.13: 1.13: 1.05: 0.83: 0.20, in female: 1.55: 1.19: 1.13: 1.00: 1.13: 0.11 (Fig. 3B). Ommatidia form a triangular structure with 11 rows.

Gnathochilarium: In the anterior portion of gnathochilarium, a distinct seta with swollen round socket is present in the stipes while the posterior part contains three rows of small setae, all with swollen round socket (peg like structure). At the base of each lingual plate five to six short setae and on both sides of the median axis, two long setae are present. Submedian transverse groove is distinct. Distal part of gnathochilarium with two rounded protruberance, inner longer with three segments while outer smaller with two segments (Fig. 3). Mandibles are joined at the base.

Mandible: External tooth long and triangular, internal tooth comparatively small, 11-13 rows of pectinate lamellae, molar plate long with a pointed horn, gnathal lobe is rectangular in shape (Fig. 4E).

Collum: Surface smooth; in male, collum broad in the middle, narrowing gently towards end, angle broadly rounded, slightly tapering at the distal (Fig. 4D). Collum of female is broader compared to that of male (2mm vs. 1.6mm) and without any tapering at the distal.

Body rings: 60 segments; in male, collum, 1^st^, preanal and anal segments are without legs; 2^nd^, 3^rd^ and 6^th^ segments with single pair of legs and rest of the segments with double pair of legs; 6^th^ segment bears gonopod; in female, 6^th^ segment with double pairs of legs. In width, head<collum<segment; 5^th^ segment in male is broadest, approximately 1.5 times than the preceding segment in width and forms a hump. Dorsum of metazonite is simple without any striation, suture distinct and prozonite with usual encircling striae (Fig. 4). Pores are small, beginning in 6^th^ segment. Anal segment with a gradually tapering long acute tail curved downwards. The tail extended 3/4^th^ of anal segment downwards (Figs. 2, 4). Marginal thickening of the anal valve is simple. The paraprocts are evenly convex and meet at a plane surface with a low rim (Fig. 4). According to classification as proposed by Hoffman (2011), it is marginate type of paraproct. Repugnatorial gland blackish, funnel shaped sac with a distinct duct located just above the midline of the body (Figs. 2, 4).

**Fig. 2.**
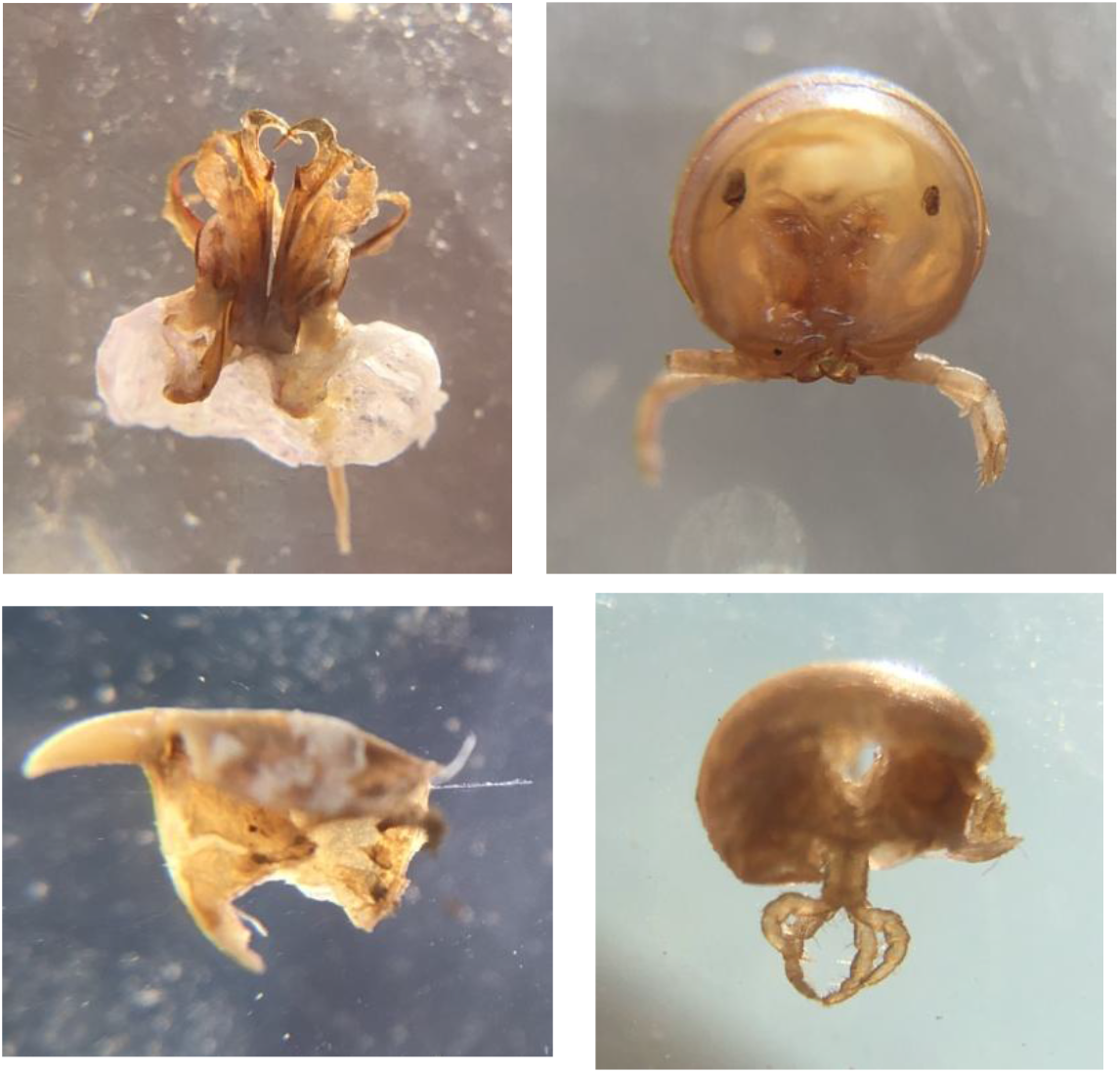
*Carlogonus bengalensis* sp. nov. (left to right, clockwise). Gonopod of male, Body ring with distinct repugnatorial gland, Tail, 2^nd^ leg.

**Fig. 3.**
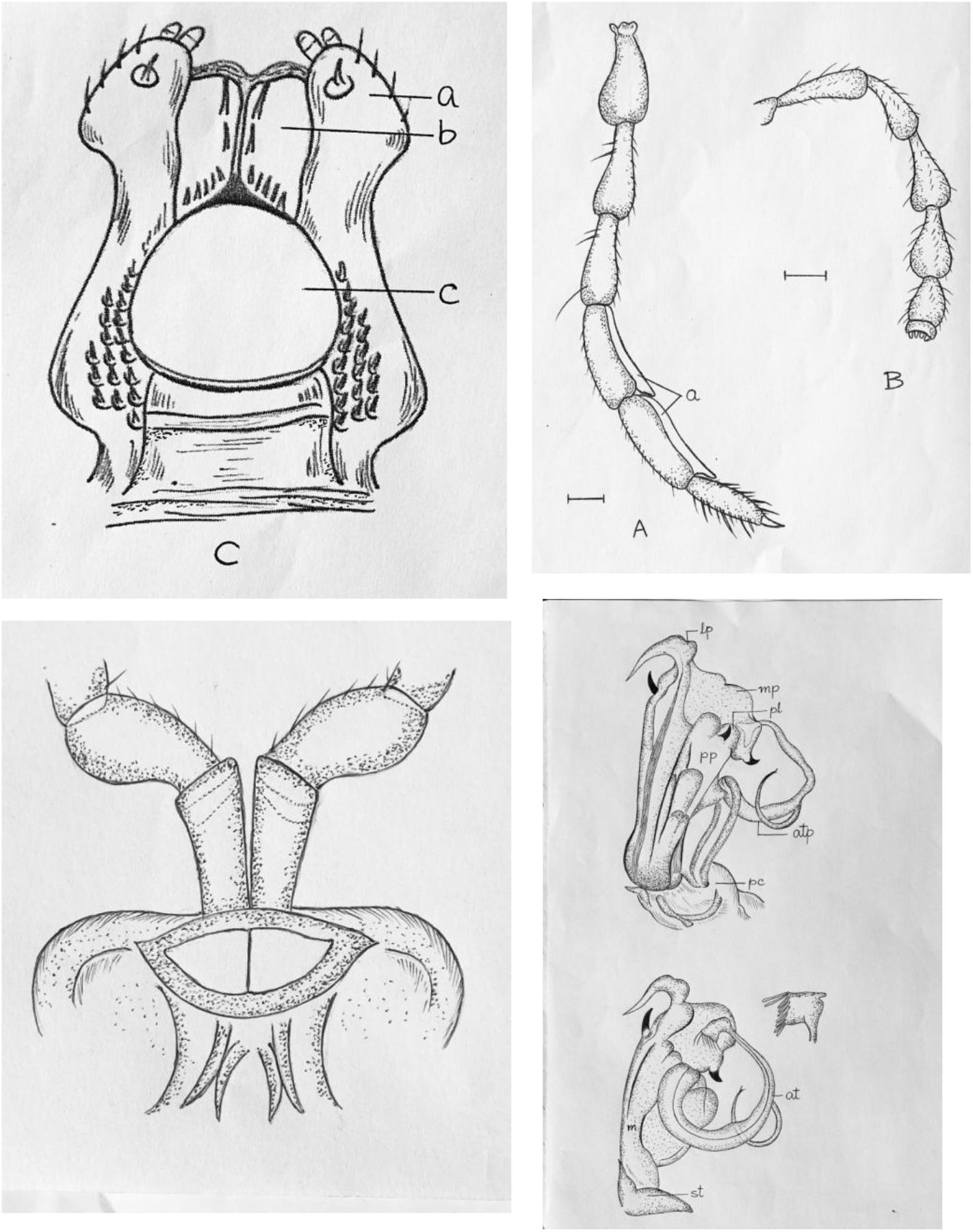
Different body parts of *Carlogonus bengalensis* sp. nov. (left to right, clockwise). Gnathochilarium: a= stipes, b= lingual plate, c= mentum; Leg (a= pad) and antenna of male; Gonopod: lp= lateral process, mp= metaplical fold of coxa, pl= proplical lobe, pp= proplical fold, atp= telopodite with antetorsal process, pc= precoxite, m= metaplica, st= sternite; 2^nd^ segment showing coxite and prefemur.

**Fig. 4.**
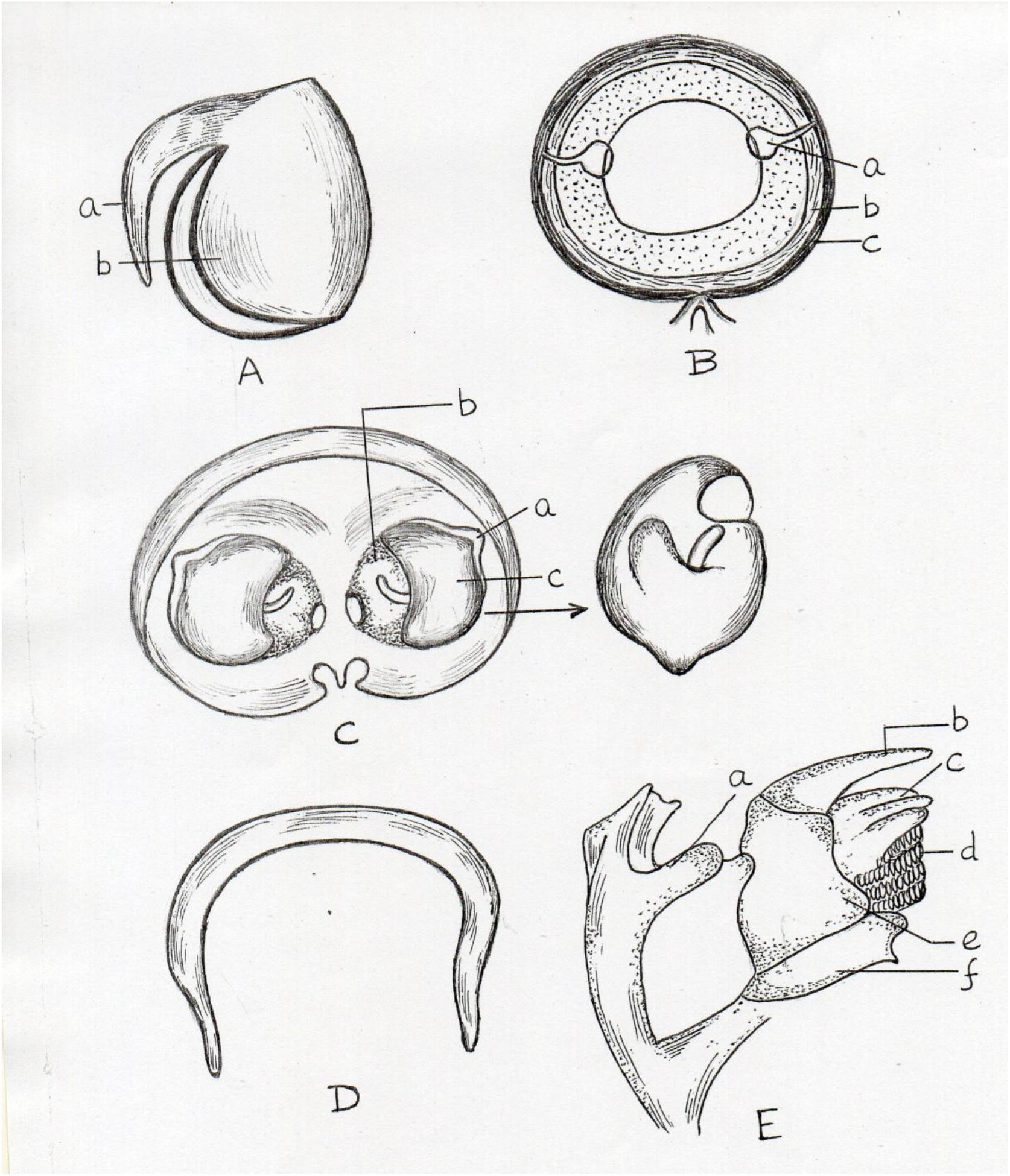
*Carlogonus bengalensis* sp. nov. A. Last segment: a= tail, b= paraproct; B. Mid body segment: a= repugnatorial gland, b= prozonite, c= metazonite; C. Ring II of female: a= vulval sac, b= operculum, c= bursa; D. Collum of male; E. Mandible: a= distal lobe of basomere, b= external tooth, c= internal tooth, d= pectinate lamellae, e= gnathal lobe, f= molar plate.

Legs: In male: leg is 0.78 of body diameter; tibia is the longest and pretarsus or nail is the smallest part; length ratio of leg parts: 1.0: 1.03: 1.17: 1.20: 1.27: 0.83: 0.27. In sternite of 2^nd^ leg of male, a curved transverse band coalesced with tracheal stalks for support. Expanded bases of the coxae not coalesced with prefemur (Figs. 2, 3).Tarsus bears numerous setae, both in the outer and inner margins, beside pretarsus, there are other seven longer secondary setae. In other parts of leg, very small outer setae are evenly distributed along the border of the outer part; prefemur and femur also have a long thread like seta in the inner portion. Whitish pads are present on post femur and tibia; in each segment pad become broader towards the end of the segment and in both segment, distal end of the pad is projected like a tongue beyond the corresponding segment (Fig. 3). These leg-pads are reduced towards the posterior end of the body and totally absent in the last leg pair. In female: leg is without pad and shorter than male; each part of leg bears a long seta towards the distal end at the inner side; length ratio of leg parts: 1.07: 1.07: 1.07: 1.07: 1.00: 1.25: 0.18; first four parts are equal in length and tarsus is the longest part; podomeres of leg attached with the vulval segment are of different lengths.

Gonopod: Gonopod is on 6^th^ segment and yellowish brown in colour. Lateral margin of proplica (anterior coxal fold) without any process; disto lateral process of proplica long, broad and hook like with a hump. Mesal process of proplica is with a red spine. Proplical fold is flat and cylindrical lobe with a curved spine which is orange in colour. Metaplica (posterior coxal fold) is broad with distally wide metaplical fold that accommodate the telopodite. A number of testaceous round spot or sigilla is present on the inner surface of the metaplical fold of which three are large and a few are small in diameter. Metaplica broad with anterior wide metaplical fold of coxa. A few testaceous round spot or sigilla is present on the inner surface of the metaplical fold of which three are large in diameter. Telopodite is long band like and curved in a circle. Proximal half of it is broad while distal half narrowing gently and end with a spatula like palette. Antetorsal process (tibial spine) occurs at about midlength of the telopodite, long, slender and almost round in shape with a bifurcated tip. Palette is wide, flat and rectangular in shape, with a horn like crest and a few blepharochatae at the broader margin. Among blepharochatae two are longer and rests are of shorter (Figs. 2, 3).

Vulva: In female 2^nd^ segment bears vulva which is slightly oval in shape and within a vulval sac, with a pointed end. Bursa of vulva is brownish but the operculum is transparent and with a curved rod-like ampulla (Fig. 4C).

Colour: Body blackish brown, leg coffee brown but the downward curved tail is yellowish. Gonopod is yellowish brown with two coloured spines (red and orange).

#### Diagnosis

Round body with 60 segments, 55mm in length, 4mm in width, 5 ^th^ segment form hump, collum of male and female are different in shape, antennomeres and podomeres lengths are different in both sexes, repugnatorial gland black and funnel shaped, marginal thickening of anal valve simple, tail curved downwards and yellow in colour, telopodite of gonopod long, flat and band like with a single antetorsal process, mesal margin of proplica bears a red spine, proplical lobe with a curved spine of orange in colour. A black spine on the lateral side of proplical fold, ante torsal process band like and curved in a circle, palette spatula like with a distinct crest and a few blepharochatae at the apical margin. Sigilla is present on the inner surface of metaplical fold.

#### Comparison

Recently Sankaran and Sebastian (2020) proposed two species groups for the 10 nominal *Carlogonus* species on the basis of a single character of the gonopod telopodite viz. “ *exaratus* group” and “*acifer* group”. As the present species bears a single tibial spine on telopodite, it belongs to “*exaratus* group”. This group includes six species – *Carlogonus exaratus* (Attems, 1936), *Carlogonus auriculus* Demange, 1983, *C. robustior, C. subvalidus, Carlogonus gayathri* Sankaran and Sebastian, 2020 and *C. bengalensis* sp. nov.

*Carlogonus bengalensis* is distinguished from *C. auriculus* by the presence of downwardly directed anterior process of anterior coxal fold vs. upwardly directed anterior process and by the absence of lateral and mesal triangular expansion on distal end of telopodite vs. presence of triangular expansion. In the present species, anterior spine like outgrowth of lateral margin of anterior coxal fold is short while it is very long in *C. robustior*. Moreover, in the later species tibial spine is short and curved but it is long and almost round in shape in the former. *C. bengalensis* differs from *C. subvalidus* by the presence of rectangular flat palette with a few blepharochaetae (vs. cylindrical with numerous blepharochaetae) and curved almost circular telopodite (vs. spiral telopodite).The present species differs from the recently discovered *C. gayathri* on several characters viz. 60 body ring (vs. 65 body ring), body colour brownish black throughout (vs. basolateral yellowish patch), long tibial spine (vs. short), expanded 5^th^ body ring in male form a hump (vs. without hump), antero lateral process of anterior coxal fold is long, wide and hook like (vs. short hook like), palette with few blepharochaetae (vs. numerous blepharochaetae). *C. bengalensis* is most nearer to *C. exaratus* as both have spine like outgrowth on the lateral margin of anterior coxal fold, mesal margin of telopodite without conical process and telopodite lacks distal, lateral and mesal triangular expansions. But it is distinguished from *C. exaratus* by the presence of long spinous outgrowth on the lateral margin of anterior coxal fold (vs. short), lateral process of posterior coxal fold with a distinct short spine (without spine), apex of the posterior coxal leaf with a short curved spine (vs. without any spine), antero mesal hook-like process of anterior coxal fold is large and wide (vs. small and narrow).

#### Natural history

According to the present knowledge, *Carlogonus bengalensis* sp. nov. is distributed only in a particular location at Suri (Dist. Birbhum, West Bengal, India). The species is observed on moist soil covered with decomposed leaves and debris. It is found during the onset of rainy season i. e. in the months of June – July. The species is nocturnal in habit and can be seen roaming in the open ground before sunrise. At this locality, other dominant millipede species are *Chondromorpha kelaarti* and *Trigoniulus lumbricinus*.

## Acknowledgement

I am deeply indebted to Late Dr. R. L. Hoffman, Virginia Museum of Natural History, USA, for identification of the millipede at the genus level. I am grateful to my son Dr. Soumendranath Bhakat and my wife for their constant inspiration and support in this phase of crisis and to my colleagues for their cooperation.

